# Frontal theta and beta oscillations during lower-limb movement in Parkinson’s disease

**DOI:** 10.1101/634808

**Authors:** Arun Singh, Rachel C. Cole, Arturo I. Espinoza, Darin Brown, James F. Cavanagh, Nandakumar Narayanan

**Affiliations:** Neurology Department, University of Iowa, Iowa City, IA, USA; Psychology Department, University of New Mexico, Albuquerque, NM, USA

**Keywords:** Parkinson’s disease, lower-limb movement, freezing of gait, oscillations, frontal region.

## Abstract

**Background:** Motor and cognitive dysfunction has been linked in patients with Parkinson’s disease (PD). EEG theta and beta rhythms are reliably associated with cognitive and motor functions, respectively. We tested the hypothesis that PD patients with lower-limb abnormalities would exhibit abnormal beta and theta rhythms in the mid-frontal region during action initiation.

**Methods:** We recruited thirty-nine subjects, including PD patients with FOG (PDFOG+; n=13) and without FOG (PDFOG−; n=13), and demographically-matched healthy subjects (n=13). Scalp electroencephalogram (EEG) signals were collected during a lower-limb pedaling motor task, which required intentional initiation and stopping of a motor movement.

**Results:** FOG scores were correlated with disease severity and cognition. PDFOG+ patients pedaled with reduced speed and decreased acceleration compared to PDFOG− patients and to controls. PDFOG+ patients exhibited attenuated theta-band (4-8 Hz) power and increased beta-band (13-30 Hz) power at mid-frontal electrode Cz during pedaling. Frontal theta- and beta-band oscillations also correlated with lower-limb movement in PD patients.

**Conclusions:** Frontal theta and beta oscillations are predictors of lower-limb motor symptoms in PD. These data provide insight into the mechanism of lower-limb dysfunction in PD, and could be used to design neuromodulation for PD-related lower-limb abnormalities.

## Introduction

Lower-body symptoms of PD include impairment in gait initiation and execution, balance, and postural instability. These symptoms are important because they can severely limit mobility and contribute to falls. Freezing of gait (FOG) is one of the most challenging lower-body symptoms in PD, and is more likely to develop as the disease progresses.^1,2^ Treatments such as levodopa and deep brain stimulation (DBS) do not reliably improve lower-body symptoms in PD.^3–5^ Lower-limb motor processes are of particular interest in PD because they combine motor and cognitive components of movement execution and movement preparation, such as anticipatory postural adjustments.^6^ However, the neuronal mechanisms and determinants of lower-limb abnormalities at the cortical level in relation to cognitive control remain largely unknown. Understanding these mechanisms is of particular important to decrease morbidity and mortality from events such as falls, and improve quality-of-life related to restricted mobility. Furthermore, elucidating mechanisms that contribute to lower-body symptoms in PD might help develop new diagnostic and therapeutic approaches.

FOG occurs during gait initiation, which is the shift from standing to walking.^7,8^ Gait initiation disturbances include delayed release of anticipatory postural adjustments and slowed movement initiation in PD.^9,10^ Previous studies have observed that reduced gait initiation time and lower-body symptoms may be related to physiological alterations in both motor and cognitive systems.^11–13^ A recent study completed in more than one hundred PD patients proposed that gait, rather than cognition, predicts decline in cognitive domains early in PD,^14^ suggesting gait might be a clinical biomarker for PD cognitive decline in early stages of the disease.

Several studies have suggested a strong connection between frontal lobe dysfunction and executive dysfunction to FOG.^15,16^ Neuroimaging studies have shown prefrontal recruitment is attenuated in PD patients with FOG,^17,18^ with the medial frontal areas accounting for motor initiation deficits in PD patients.^19^ Medial frontal regions such as supplementary motor area, leg motor cortex, and anterior cingulate cortices interact to instantiate motor and cognitive control.^20–22^

Medial frontal scalp EEG can identify neurophysiological signatures of both motor and cognitive control. Beta rhythms between 13-25 Hz are associated with motor control and increase with upper-limb bradykinesia and upper-limb motor impairments in PD.^23,24^ This line of evidence predicts that increased frontal beta activity underlies impaired motor initiation in FOG. Theta rhythms between 4-8 Hz are associated with cognitive control are attenuated in PD.^20,22,25,26^ Because PD patients with FOG have greater deficits in executive function and attention compared to PD patients without FOG,^27^ we hypothesized that attenuated theta activity might also underlie FOG in PD.

We tested these hypotheses by investigating the relationship of FOG in PD by collecting cortical EEG during a pedaling task. We examined the relationship of these neural signatures with cognitive function and motor symptoms severity of PD. We found attenuated mid-frontal theta activity as well as increased frontal beta activity in PD patients with FOG. These findings provide insight into the neuronal mechanisms of lower-limb symptoms of PD. Because these frontal rhythms engage cortical and subcortical neuronal populations involved in motor and cognitive control and can be targeted by neuromodulation techniques,^20,28–30^ these data may contribute to novel therapies that improve FOG in PD.

## Methods

### Subjects and Clinical Assessments

We recruited thirty-nine subjects (13 PD patients with FOG, “PDFOG+”; 13 PD patients without FOG, “PDFOG–”, and 13 demographically-matched healthy subjects, “controls”). All protocols were approved by the University of Iowa Office of the Institutional Review Board (IRB). All PD patients met UK Parkinson’s Disease Brain Bank Criteria^31^ for the diagnosis of idiopathic PD and were recruited in the Department of Neurology and Movement Disorders at the University of Iowa Hospitals and Clinics. All subjects provided written informed consent. Subjects were paid $30/h for participation. All PD patients were treated with levodopa medication. Levodopa-equivalent dosages (LED) for each patient are shown in Table 1.

**TABLE 1.** Demographic, disease, motor, and cognitive characteristics

|  | **Control**<br>**(N = 13)** | **PDFOG–**<br>**(N = 13)** | **PDFOG+**<br>**(N = 13)** | ***P* Value** |
| --- | --- | --- | --- | --- |
| Gender, M/F | 8/5 | 9/4 | 8/5 |  |
| Age, years | 69 (2.3) | 65.2 (2.4) | 69.5 (2.6) | >0.05<sup>a</sup> |
| Disease Duration, years | - | 4.8 (1.0) | 7.5 (1.0) | 0.06 |
| LED, mg/day | - | 682.7 (99.1) | 1158.8 (134.4) | 0.01* |
| **Cognition Characteristics** |  |  |  |  |
| MOCA (0-30) | 26.6 (0.53) | 24.8 (0.8) | 21.8 (1.2) | <0.01<sup>b**</sup> |
| **Motor Characteristics** |  |  |  |  |
| UPDRS III (0-56) | - | 10.5 (1.7) | 19.2 (1.3) | <0.01** |
| H & Y stage, on (1-5) | - | 1.6 (0.2) | 2.4 (0.3) | 0.03* |
| **Freezing of Gait (FOG)** |  |  |  |  |
| FOG questionnaire (0-24) | - | 3.7 (0.9) | 14.8 (0.8) | <0.01** |

Values were expressed as mean (standard error of mean).
<sup>a</sup>One-way ANOVA was applied. Two-sample *t* test was used for comparison between two groups. <sup>b</sup>PDFOG+ vs. control subjects. \*Groups significantly different at $P < 0.05$ . \*\*Groups significantly different at $P < 0.01$ . Abbreviations: Male, M; Female, F; Montreal Cognitive Assessment, MOCA; motor Unified Parkinson's Disease Rating Scale, UPDRS III; Hoehn & Yahr scale, H & Y.

Patients participated in research for one 4-hour session after a clinic appointment. The visit started with questionnaires, including the Freezing of Gait Questionnaire (FGQ)^32^ to evaluate gait impairments in patients. Participants then completed multiple cognitive tasks and the pedaling motor task during EEG recording. At the end of the EEG session we administered the Montreal Cognitive Assessment (MOCA) to measure cognition^33^ and the motor Unified Parkinson’s Disease Rating Scale (UPDRS III)^34^ to measure motor symptoms of PD. Patients with a FOG score > 10 and a FGQ item 3 score > 0 were classified as PDFOG+. Table 1 shows the subject’s demographic and clinical information in detail.

We performed non-parametric Spearman correlation analyses between motor (FOG and UPDRS III) and cognitive assessment (MOCA) scores of PD patients. We further performed partial correlation to determine the relationship between disease duration and FOG when controlling for MOCA, and between disease duration and MOCA when controlling for FOG.

### Lower-limb Motor Task and Analysis

We used a pedaling motor task (Fig. 1A) to study lower-limb movement control for the following reasons: (1) to minimize fall-risks in PD patients with marked gait abnormalities, (2) pedaling generates EEG signals with minimal movement artifact, (3) pedaling can requires bilateral coordination, and (4) pedaling kinematics analysis allow for detailed measures of lower-limb function. PDFOG+ patients can experience discontinuous changes in speed during continuous pedaling related to FOG.^35,36^ Patients were seated on a chair for the pedaling task (2 blocks of 30 trials). For each trial, a GO cue (green circle) appeared on the stimulus presentation computer and the subject was instructed to complete one rotation. They stopped the pedals in the starting position and waited for the next GO cue (3 seconds inter-trial interval).

**FIG. 1.**
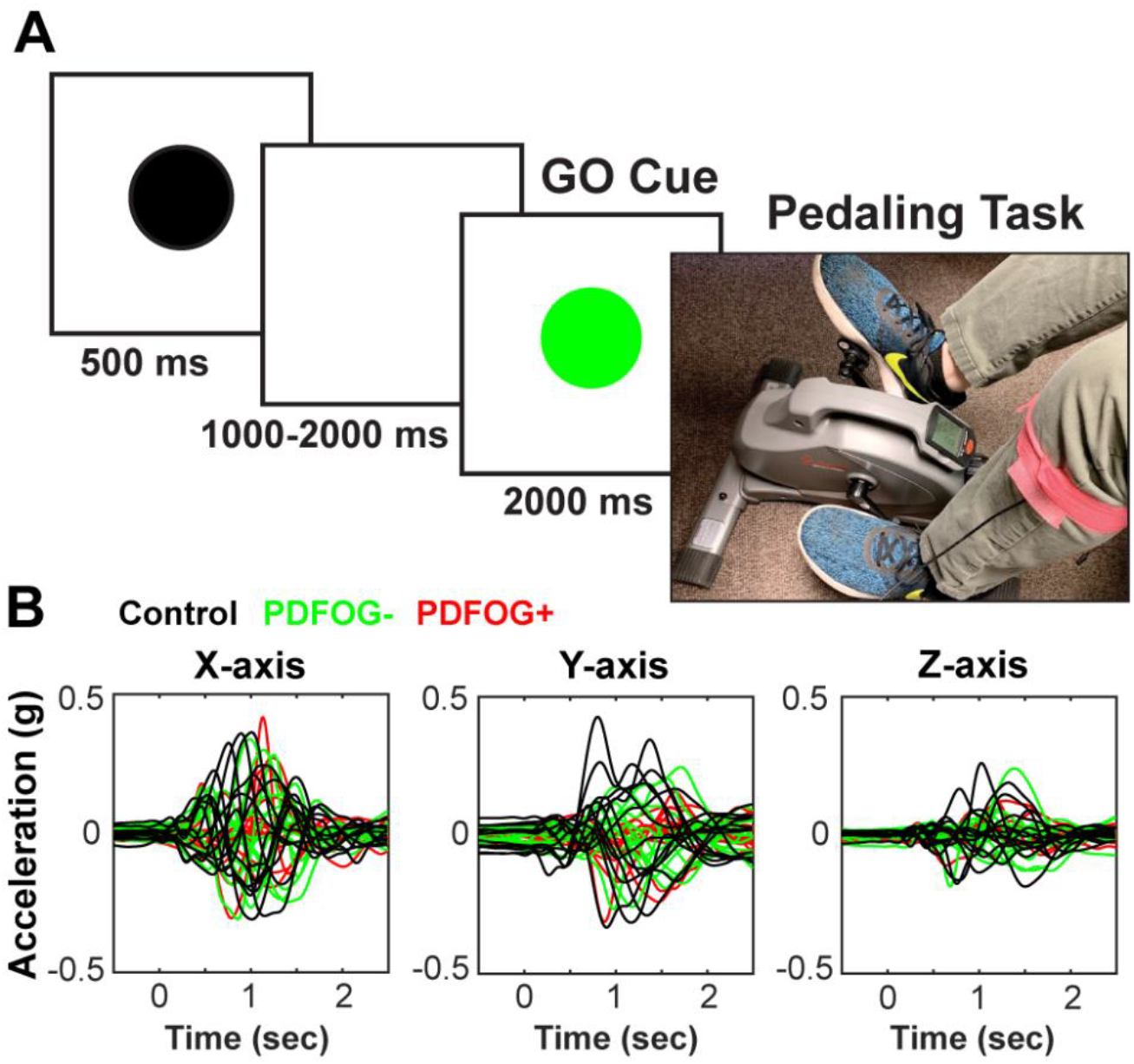
Experimental design. (A) In this task, a black “warning” cue appeared to alert the subject to pay attention. Within 1000-2000 ms, a green GO cue instructed subject to complete one rotation of the pedals. (B) Accelerometer signals were collected from tri-axial accelerometer and segmented from the GO-cue (−500 to 2500 ms) and averaged to plot the mean trace of X-, Y-, and Z-axes, in control, PDFOG–, and PDFOG+ participants. FOG: Freezing of Gait; PDFOG–: PD patients without FOG; PDFOG+: PD patients with FOG.

We computed the linear speed for each pedaling trial from 3-axis accelerometer signals. We selected 0-2000 ms from the GO cue as the time-window to analyze speed and acceleration because participants took >2000 ms to complete one pedal rotation (Fig. 1B). Accelerometer signals (X-, Y-, and Z-axes) were detrended and low-pass filtered at 5 Hz. Mean speed was computed for each axis for each trial, and then averaged across axes. Finally, we averaged all trials to represent the final mean speed (g*s) of the pedaling task for each subject. Additionally, to compute the time to achieve maximum acceleration, we picked the accelerometer signal (g) in which we observed highest acceleration using the Matlab “findpeaks” function. We averaged the time to reach peak acceleration across all trials to represent the mean time (peak time) for each subject.

For comparisons of speed and maximum acceleration time between control, PDFOG–, and PDFOG+, we performed one-way analyses-of-variance (ANOVA). We used two-tailed t-tests with an alpha level of 0.05 to compare between two groups. Furthermore, we also performed Spearman correlation analyses between kinematic parameters (pedaling speed and maximum acceleration time) and motor (FOG and UPDRS III) and MOCA scores of PD patients.

### EEG Recording and Analysis

Electroencephalogram (EEG) signals were collected during the lower-limb pedaling motor task from a customized 64-channel cap (Easycap.Inc) using high-pass filter of 0.1 Hz with a sampling rate of 500 Hz (Brain Products). Online reference and ground channels were Pz and FPz, respectively. Data were epoched around the GO stimulus onset (−1000 to 3000 ms). Data were then re-referenced to an average reference. FP1, FP2, FT10, TP9, and TP10 channels were removed, as they tend to be influenced by eyeblink / muscle artifact, leaving 59 electrodes for analysis. Of note, we used a specialized cap without FT9. Bad channels and epochs were identified using a conjunction of the FASTER algorithm^37^ and pop_rejchan from EEGlab and were subsequently interpolated and rejected respectively. Eye blinks were removed following ICA.

All analyses in the current report are from the mid-frontal Cz vertex electrode. After preprocessing, time-frequency measures were computed by complex Morlet wavelets,^38^ also explained in our previous report.^20^ For time-frequency computation, each epoch was cut in length (−500 to +2000 ms), frequency bands between 1-50 Hz in logarithmically-spaced bins were selected, and power was normalized by conversion to a decibel (dB) scale. The baseline for each frequency consisted of the average power from −300 to −200 ms prior to the onset of the GO stimuli. This short baseline period is common in the field since a small-time sample reflects the wavelet-weighted influence of longer time and frequency periods. We restricted our analyses to electrode Cz and a-priori time-frequency Region of Interest (tf-ROI). We exported mean power values in the theta-band (3.5-7.5 Hz) and beta-band (12.5-30 Hz) from 0-2000 ms following the GO cue for statistical comparisons. Additionally, we exported theta- and beta-bands power values in 0-400 ms (time of motor initiation) range.

ROIs were preselected for the theta- and beta-bands. To quantify time-frequency analyses, we set the size threshold for chance occurrence of the statistical cluster against 1000 permutations of group labels as the one-dimensional cluster mass at the 95th percentile. We used linear regression to analyze the difference in power for each tf-ROI between control, PDFOG–, and PDFOG+, while controlling for MOCA. We selected the control group to be the reference group, and we first compared both PDFOG+ and PDFOG– to the reference group. Linear regression was done in R 3.5.1, with the package “stats”. Pairwise comparisons were evaluated by fitting an additional linear model with only two groups, while covarying for MOCA. All pairwise comparisons were tested with an alpha level of <0.05. Finally, we used Spearman correlation analyses to evaluate the relationships between power values during gait initiation (theta- and beta-bands; 0-400 ms) and motor and cognitive characteristics (FOG, MOCA and UPDRS III) of PD patients.

## Results

### Patient demographics

Table 1 shows the group comparisons for all motor, cognitive, and other clinical characteristics. No differences were found for age between PD and control groups. MOCA scores were significantly different between PD patients and control subjects, with PD patients scoring lower than control subjects (*P*=0.01). PDFOG+ patients had longer disease duration and significantly higher LED (*P*=0.01) than PDFOG– patients. Subsequent group analyses will control for MOCA and disease duration. PDFOG+ patients had higher UPDRS III, higher FOG, and lower cognitive (MOCA) scores than PDFOG– patients (see Table 1).

### Motor and Cognitive Assessment Scores

Previous reports have suggested that FOG can be related to motor and cognitive impairments.^39,40^ Accordingly, in PD patients, we observed a positive correlation between FOG and motor function as measured by UPDRS III (Fig.2A; *P*=0.001), and a negative correlation between FOG and cognition as measured by MOCA (Fig. 2B; *P*=0.02). We observed a positive correlation between FOG scores and disease duration (Fig. 2C; *P*=0.01), even when controlling for MOCA (*P*=0.03). Of note, there was no significant negative correlation between disease duration and MOCA scores (Fig. 2C; *P*=0.08). These data suggest that lower-limb movements tend to be worse with longer disease duration, while cognition is not related to duration.^14^

**FIG. 2.**
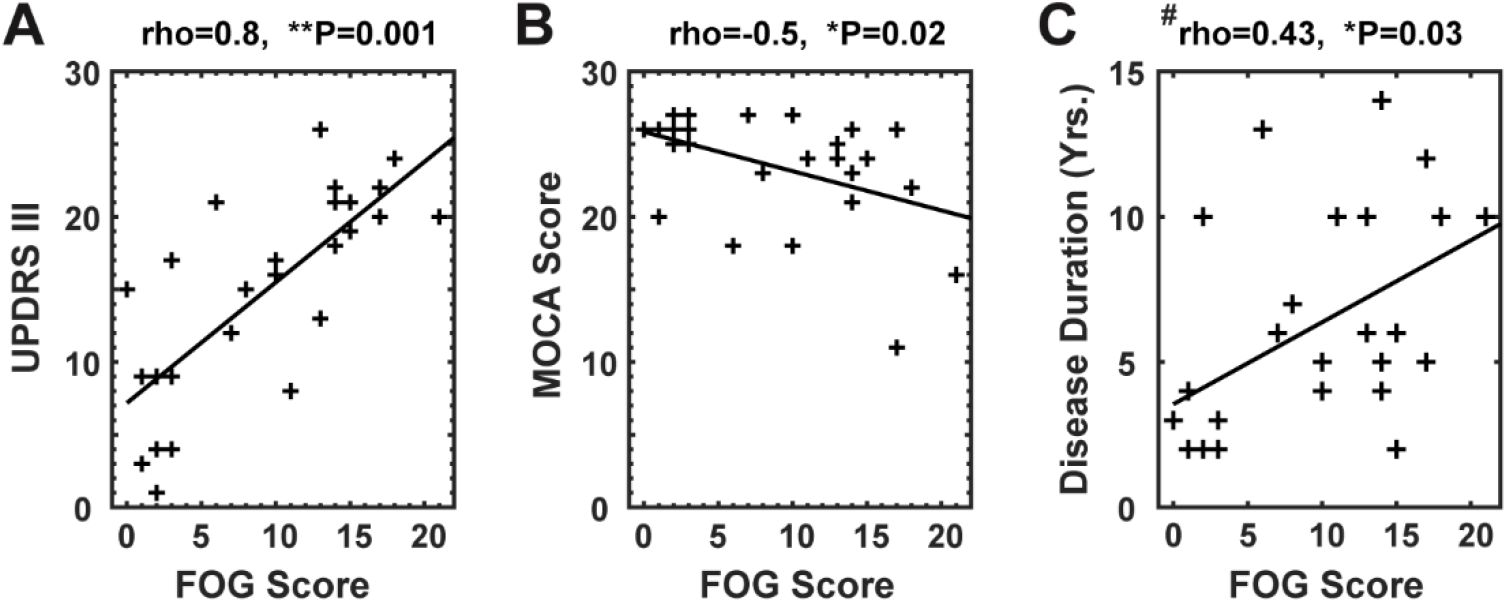
Relationship between FOG and motor /cognitive characteristics of PD. (A-B) FOG questionnaire scores as a measurement of lower-extremity impairment showed a significant positive correlation with UPDRS III scores and a significant negative correlation with MOCA scores. (C) ^#^Partial correlation analysis showed a significant correlation between disease duration (DD) and FOG, but not DD and MOCA scores. *Significance correlation level <0.05; **Significance correlation level <0.01. rho = correlation coefficients; UPDRS III: motor Unified Parkinson’s Disease Rating Scale; FOG: Freezing of Gait; MOCA: Montreal Cognitive Assessment.

### Kinematic Results of Pedaling Task

PD patients are impaired in lower-limb movements as measured by the speed of motor task initiation and execution. Our results showed that all PD patients (both PDFOG– and PDFOG+) executed the lower-limb pedaling motor task with significantly lower speed compared to control subjects (Fig. 3A; F(2,36)=9.9, *P*=0.001, η_p_^2^=0.35; PDFOG+ vs. controls, a decrease of 57% *P*=0.001; PDFOG– vs. controls, a decrease of 40%, *P*=0.01). Correlation results revealed relationships between pedaling speed and FOG scores (Fig. 3A top; P=0.01) and UPDRS III scores (Fig. 3A middle; *P*=0.001), but not between pedaling speed and MOCA scores (Fig. 3A bottom; *P*=0.3).

**FIG. 3.**
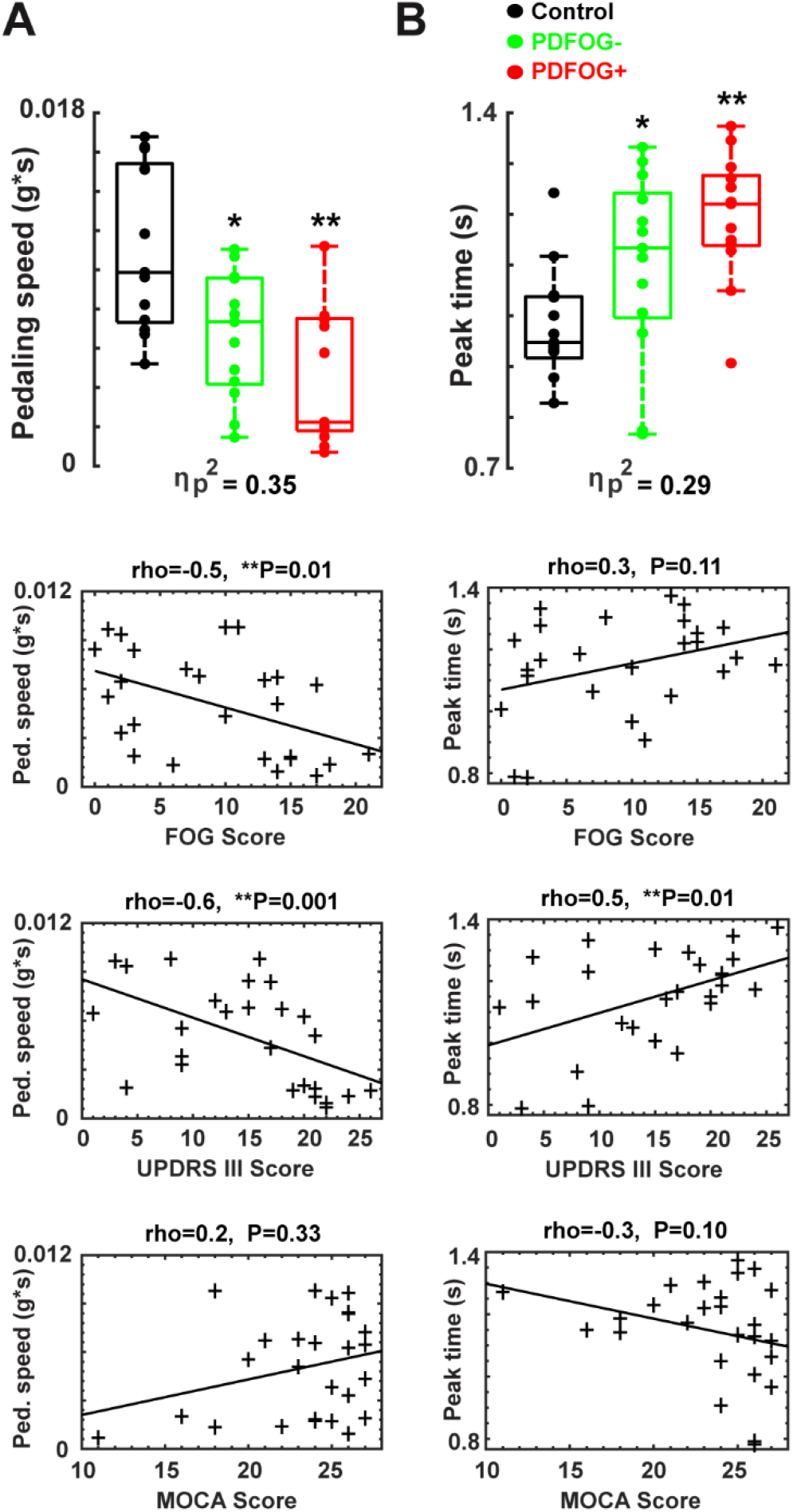
Pedaling kinematics. (A) Box plots show that mean pedaling linear speed was the lowest in PDFOG+, and significantly correlated with FOG scores, and UPDRS III scores, but not with MOCA scores. (B) Box plots also show that PDFOG+ took the most time to reach maximum acceleration (peak time) during the pedaling task, and a significant correlation was found between peak time and UPDRS III scores only, but not FOG scores and MOCA scores. \**P*<0.05 vs control subjects; \*\**P* <0.01 vs control subjects; *Significance correlation level <0.05; **Significance correlation level <0.01. Effect size was symbolized by partial eta-squared (η_p_^2^). rho = correlation coefficients; FOG: Freezing of Gait; UPDRS III: motor Unified Parkinson’s Disease Rating Scale; MOCA: Montreal Cognitive Assessment. PDFOG+: PD patients with FOG; PDFOG–: PD patients without FOG.

Additionally, we also computed each subjects’ time to achieve maximum acceleration and found that all PD patients took more time to reach peak acceleration compared to control subjects (Fig. 3B; F(2,36)=7.7, *P*=0.002, η_p_^2^=0.29; PDFOG+ vs. controls, an increase of 24% *P*=0.001; PDFOG– vs. controls, an increase of 14% *P*=0.04). PDFOG+ also took more time to reach peak acceleration compared to PDFOG– (Fig. 3B; an increase of 8% *P*=0.14). Time to reach peak acceleration was significantly correlated with UPDRS III scores (Fig. 3B middle; *P*=0.01) but not FOG scores (Fig. 3B top; *P*=0.1) or MOCA scores (Fig. 3B bottom; *P*=0.09). These results indicate that PD patients with more severe motor symptoms exhibited greater slowing on the lower-limb motor task.

### Pedaling Task-related Power Changes at Frontal Region

We tested the hypothesis that PDFOG+ patients had attenuated mid-frontal theta activity and increased beta activity during lower-limb motor task. In support of this idea, the tf-ROI theta-band showed significantly attenuated power in PDFOG+ patients compared to both controls and PDFOG– at the time of pedaling initiation and during the execution of the pedaling task (Fig. 4A and B).

**FIG. 4.**
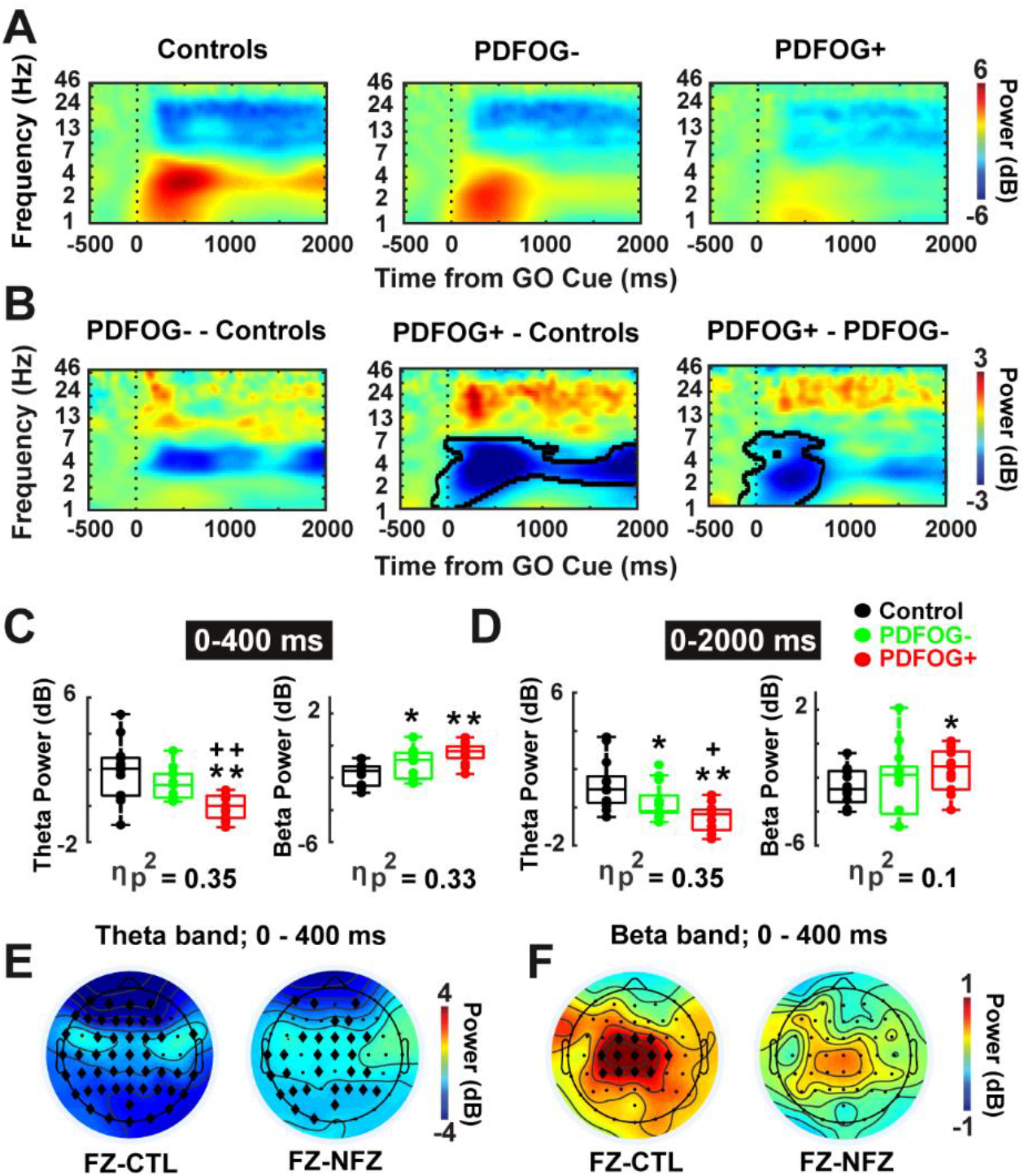
FOG is associated with attenuated theta-band and amplified beta-band power in PD. (A-B) Time-frequency analysis showed reduced theta and amplified beta power at frontal scalp electrode (vertex or “Cz”) during pedaling task in PDFOG+ patients as compared to PDFOG– patients and control subjects. PDFOG– patients showed low theta power and high beta power compared to control subjects. (C) Box plots displayed the power values from two tf-ROIs (theta and beta power values at 0-400 ms and 0-2000 ms time-window) during pedaling task. (E-F) Topography plots (PDFOG+ patients versus controls and PDFOG+ patients versus PDFOG– patients) indicated reduced theta and increased beta activity at the frontal region in PDFOG+ patients. B: Permutation-corrected statistical significance *P*<0.05 outlined in bold lines. \**P*<0.05 vs control subjects; \*\**P*<0.01 vs control subjects; ^+^*P*<0.05 vs PDFOG–; ^++^*P*<0.01 vs PDFOG–; Effect size was symbolized by partial eta-squared (η_p_^2^); diamonds show significant electrodes in E and F. FOG: Freezing of Gait; PDFOG+: PD patients with FOG; PDFOG–: PD patients without FOG. CTL: control subjects; NFZ: PDFOG–; FZ: PDFOG+.

Notably, PDFOG+ and PDFOG– patients had distinct MOCA scores. To control for these differences, we turned to separate linear regression models for theta- and beta-bands power tf-ROIs, with each tf-ROI as the outcome variable and group (PDFOG+, PDFOG–, and control) and MOCA as explanatory variables. Group effects will be described with reference to the mean of control subjects (i.e. dummy coding with reference group being control subjects), covarying for the continuous variables of MOCA.

Regression analyses showed a main effect of group on frontal theta-band power during the motor initiation epoch (0-400 ms; Fig. 4C left side; F(2,35)=9.6, *P*=0.001; η_p_^2^=0.35). When covarying for MOCA, frontal theta-band power in PDFOG– was not significantly different from the control group (*P*=0.09). However, theta power in PDFOG+ differed significantly both from the control group (a decrease of 101% *P*=0.001), and PDFOG– (a decrease of 102% *P*=0.002). Results also revealed a main effect of group on frontal theta power during the entire 2s window of pedaling epoch (0-2000 ms; Fig. 4D left side; F(2,35)=9.4, *P*=0.001; η_p_^2^=0.35). PDFOG+ had significantly lower theta power than PDFOG– (a decrease of 334%; *P*=0.03) and controls (a decrease of 101%; P=0.0001). PDFOG– patients even showed significant reduction in theta power (at 0-2000 ms) during pedaling compared to control subjects (a decrease of 80% *P*=0.02).

We also observed differences between groups in frontal beta-band activity during motor initiation (Fig. 4A and B). Analyses of tf-ROIs beta-band showed a main effect of group during the motor initiation epoch, while controlling for differences in MOCA (0-400 ms; Fig. 4C right side; F(2,35)=8.8, *P*=0.001; η_p_^2^=0.33). Beta power was increased in PDFOG+ patients and PDFOG– patients compared to controls (PDFOG+ vs. controls, an increase of −73% *P*=0.002 and PDFOG– vs. controls, an increase of −43% P=0.03). Beta power was also higher in PDFOG+ compared to PDFOG– but it was not significant (an increase of −54% P=0.9). We repeated these analyses for the entire 2s time window of pedaling (0-2000ms; Fig. 4D right side). Surprisingly, we did not observe differences in frontal beta-band power for the entire 2s time window (F(2,35)=1.4, P=0.3; η_p_^2^=0.1). However, beta-band power was significantly increased in PDFOG+ compared to controls (an increase of −38% *P*=0.01), but not in PDFOG– compared to controls (*P*=0.3) and PDFOG+ compared to PDFOG– (*P*=0.3).

Scalp topography plots indicated the presence of attenuated theta-band and amplified beta-band oscillations in the frontal region during movement initiation in PDFOG+ compared to control and PDFOG– (Fig. 4E and F).

Further analysis confirmed a significant relationship between FOG scores and theta-band power values during motor initiation (0-400 ms; Fig. 5A top; *P*=0.01), and between FOG scores and beta-band power values during motor initiation in PD patients (0-400 ms; Fig. 5B top; *P*=0.05). Interestingly, MOCA scores trended to correlate with theta-band power values (0-400 ms; Fig. 5A bottom; *P*=0.056) and UPDRS III scores were correlated with beta-band power values in PD patients (0-400 ms; Fig. 5B middle; *P*=0.026). These results suggest that abnormal theta-band power is related to cognitive control, and beta-band activity is related to the motor control. Overall, these results suggest that the attenuated theta-band activity and increase beta-band activity in the frontal lobe could be cortical predictors of lower-limb motor dysfunction in PD patients.

**FIG. 5.**
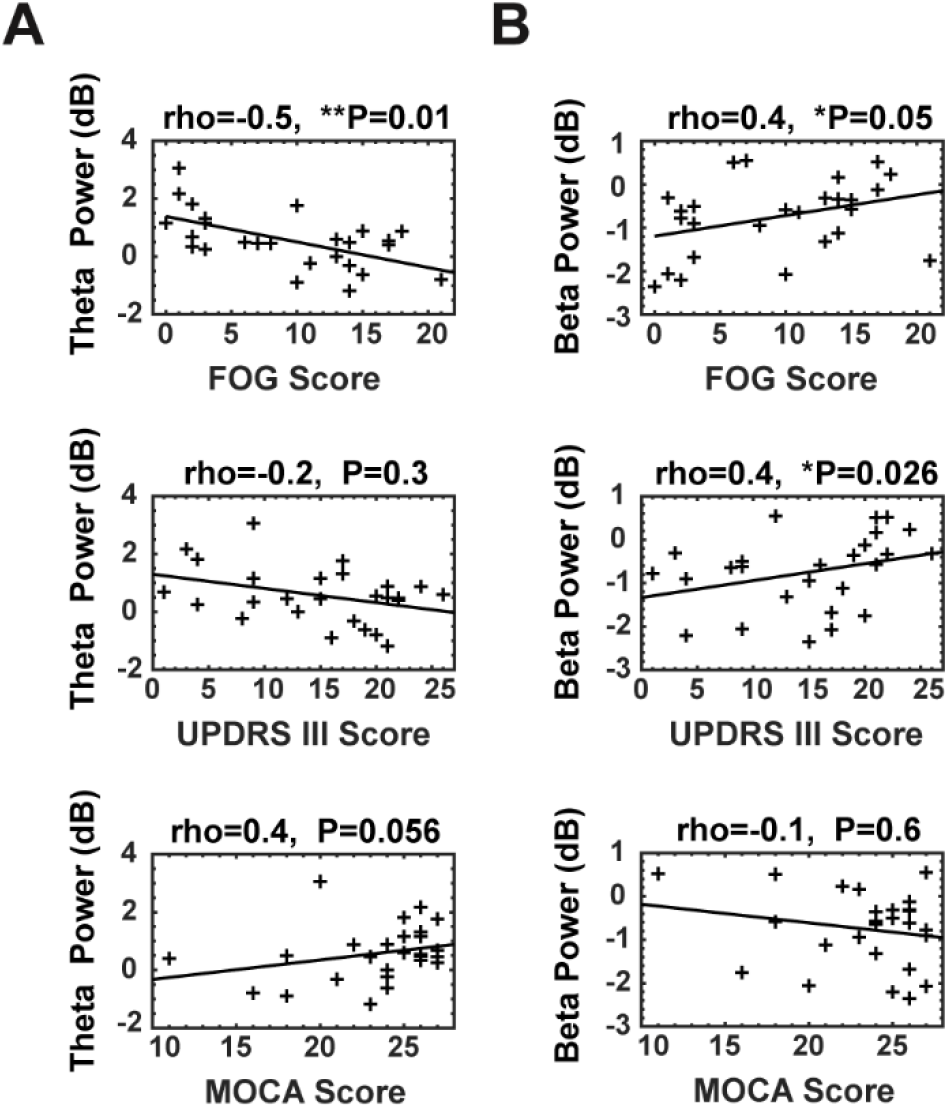
Correlation analysis between power values of theta and beta-band and motor and cognitive characteristics in PD. (A) A significant correlation was seen between theta power values and FOG scores and MOCA scores, but not with UPDRS III scores. (B) A significant correlation was observed between beta power values and FOG scores and UPDRS III scores, but not with MOCA scores. *Significance correlation level <0.05; **Significance correlation level <0.01. rho = correlation coefficients; FOG: Freezing of Gait; UPDRS III: motor Unified Parkinson’s Disease Rating Scale; MOCA: Montreal Cognitive Assessment. All power within tf-ROIs of 0-400 ms.

## Discussion

We explored the neural basis of FOG in PD patients. We hypothesized that attenuated theta- and enhanced beta-rhythms would underlie FOG in PD. Accordingly, we found evidence of decreased mid-frontal theta activity during motor initiation and pedaling, and evidence of increased beta activity particularly during motor initiation. Linear regression suggested that this impairment was independent of differences in MOCA. These data provide insight into the neural mechanisms of FOG in PD.

Mid-frontal theta-rhythms have been associated with cognitive control, and can be associated with impairments in cognitive control in PD patients.^20,25^ Our past work has suggested that frontal theta-rhythms are not modulated by levodopa.^20^ We found that PDFOG+ patients had impaired cognitive control relative to PDFOG– patients and attenuated frontal theta-rhythms. PD patients may have impaired attention to gait when walking under dual-task conditions;^41^ indeed, PD patients had marked gait variability during stable walking and performing a cognitively challenging task. Neuroimaging studies have shown correlations between FOG scores and executive impairments in PD and suggested that FOG could be associated with dysfunction within frontoparietal regions of the cortex which sub-serve cognitive and executive functions.^17,18,42^ In one human study, prefrontal volume was associated with gait such that smaller prefrontal volume was related with poorer gait performance.^43^ Further, intracortical recordings in cats demonstrated a critical role for frontal regions during gait, particularly during stepping movements.^44^ These studies imply that there might be a possible impairment in frontal regions controlling movements such that attentional and executive function networks in executing lower-limb motor performance. Currently, targeted cognitive training as a non-pharmacological intervention has been performed in PDFOG+ to reduce the severity of FOG and improve lower-limb movements.^13,45^

It has been shown that reduced theta activity in PD patients correlates with deficits in sequence acquisition and stabilization of newly acquired movement patterns.^46^ Mid-frontal theta rhythms are a mechanism of cognitive control that is impaired in PD.^20,28,47,48^ Therefore, it seems that theta oscillations may contribute to sensorimotor integration needed to execute a task.^49^

Increased beta-rhythms are associated with decreased movement in PD patients.^21,24^ Critically, levodopa attenuates beta-rhythms, and PDFOG+ patients had increased beta activity despite increased LED. These data argue that differences in beta-rhythms were not a result of differences in levodopa, although our patients were not tested during levodopa ‘OFF’ periods due to highly-increased fall risks. It has been proposed that the frontal region interacts with the basal ganglia nuclei during lower-limb motor task via beta-band oscillations.^21,24^ Beta-band synchrony within the cortico-basal ganglia circuit promotes tonic activity that slows down lower-limb movements, thus providing further evidence for the role of aberrant frontal beta oscillations in lower-extremity abnormalities.^23^ Increased frontal beta oscillations in PDFOG+ during lower-limb initiation may suggest disconnectivity in top-down neural circuitry executing movement. Frontal regions may play an important role in top-down signaling to guide adjustment of preparatory and execution plans during motor tasks, and these processes may malfunction in PD.^50^ Overall, the present data suggest an involvement of frontal theta and beta oscillations in initiation and execution of lower-extremity movements.

Similar to upper-limb movements, PD patients execute lower-limb movements with lower speed.^51–53^ Previous reports have confirmed that PDFOG+ walk with lower speed compared to PDFOG– or age-matched healthy subjects ^24,54^ and take more time to initiate movement.^9,10,55^ Here, subjects initiated the lower-limb pedaling task after seeing a visual “GO” cue. Interestingly, visual and auditory cues may affect lower-limb movement, such that focusing attention on visual cues during the task might compensate for a proprioceptive processing deficit in PD.^56,57^ Our results demonstrate that initiation time was higher in PDFOG+, and speed to execute the task was lower, but it is also possible that patients would have shown even more impairment without the visual cue.

Overall, our data suggest an imperative role of motor and cognitive determinants in performing lower-limb movement.^6,13,45,58^ Typically, lower-limb abnormalities are diagnosed as an axial motor symptom using FOG scores in advanced PD patients.^34^ Table 1 indicates the significant differences in both motor and cognitive aspects between control subjects and PD patients, and between PDFOG+ and PDFOG–. We also found that MOCA scores were different between PDFOG+ and PDFOG– and correlated with FOG scores. MOCA is used to evaluate cognitive function, including many domains such as attention, visuospatial skills, orientation, executive functions, and memory which have effect on advanced PD patients.^33^

Limitations of our study include: (1) We did not study patients during walking movements or freezing episodes. Of note, pedaling induces less movement-related artifacts and can be used in closed-loop neuromodulation experiments, and has less inherent fall risks; (2) Cortical dysfunction in PD can be complex as some PD patients may have enhanced frontal function.^59,60^ However, our analyses captured both increased and decreased patterns of frontal oscillatory activity; (3) Reduced brain activity in frontal areas is a basic abnormality in motor performance in PD,^61^ but we did not disambiguate upper vs. lower motor movements; (4) we did not study our patients OFF levodopa, although our prior work indicated that midfrontal theta abnormalities in PD do not depend on levodopa, and beta abnormalities depend on movement rather than levodopa, and we had PDFOG+ and PDFOG– perform a similar motor task; and (5) EEG signals are recorded at the scalp and are inherently limited measures of neuronal activity. We are exploring intracranial recordings, brain imaging, and experiments in animal models to further elucidate the cortical basis of theta and beta abnormalities in FOG. Basal ganglia and brain stem networks also play a key role in lower-limb movements such as gait.^23,24,62^

These results suggest that synergistic interaction of motor and cognitive control systems contributes to the execution of a lower-limb motor task. The fine initiation and execution of a lower-limb movement rely on high theta and low beta oscillatory activities in the frontal region, respectively. Increased frontal cortical theta activity might be related to stabilization of cognitive control and decreased frontal cortical activity might be related to motor control as indicated by reduced susceptibility to interference to initiate and execute lower-limb movement. Our findings could link lower-limb dysfunction to a specific motor process, namely a failure to recruit lower-limb motor control processing at key moments. We also suggest that theta and beta oscillations can be used as a feature for closed-loop neuromodulation in the future to improve lower-limb function in PD and movement disorders.

## Acknowledgements

The authors thank all participants in this study. AS, RCC, AIE, JG and NN are supported by NINDS R01100849. All data and code for this study will be available on-line at narayanan.lab.uiowa.edu or PREDiCT.

## References

1. Bloem BR, Hausdorff JM, Visser JE, Giladi N. Falls and freezing of gait in Parkinson’s disease: a review of two interconnected, episodic phenomena. Mov Disord 2004;19:871–884.

2. Boonstra TA, van der Kooij H, Munneke M, Bloem BR. Gait disorders and balance disturbances in Parkinson’s disease: clinical update and pathophysiology. Curr Opin Neurol 2008;21:461–471.

3. Rocchi L, Carlson-Kuhta P, Chiari L, Burchiel KJ, Hogarth P, Horak FB. Effects of deep brain stimulation in the subthalamic nucleus or globus pallidus internus on step initiation in Parkinson disease: laboratory investigation. J Neurosurg 2012;117:1141–1149.

4. St George RJ, Nutt JG, Burchiel KJ, Horak FB. A meta-regression of the long-term effects of deep brain stimulation on balance and gait in PD. Neurology 2010;75:1292–1299.

5. Xie T, Padmanaban M, Bloom L, et al. Effect of low versus high frequency stimulation on freezing of gait and other axial symptoms in Parkinson patients with bilateral STN DBS: a mini-review. Transl Neurodegener 2017;6:13.

6. Amboni M, Barone P, Hausdorff JM. Cognitive contributions to gait and falls: evidence and implications. Mov Disord 2013;28:1520–1533.

7. Callisaya ML, Blizzard L, Martin K, Srikanth VK. Gait initiation time is associated with the risk of multiple falls-A population-based study. Gait Posture 2016;49:19–24.

8. Henriksson M, Hirschfeld H. Physically active older adults display alterations in gait initiation. Gait Posture 2005;21:289–296.

9. Delval A, Tard C, Defebvre L. Why we should study gait initiation in Parkinson’s disease. Neurophysiol Clin 2014;44:69–76.

10. Rosin R, Topka H, Dichgans J. Gait initiation in Parkinson’s disease. Mov Disord 1997;12:682–690.

11. Killane I, Fearon C, Newman L, et al. Dual Motor-Cognitive Virtual Reality Training Impacts Dual-Task Performance in Freezing of Gait. IEEE J Biomed Health Inform 2015;19:1855–1861.

12. Martin K, Blizzard L, Garry M, Thomson R, McGinley J, Srikanth V. Gait initiation in older people--Time to first lateral movement may be the measure of choice. Gait Posture 2011;34:374–378.

13. Walton CC, Mowszowski L, Gilat M, et al. Cognitive training for freezing of gait in Parkinson’s disease: a randomized controlled trial. NPJ Parkinsons Dis 2018;4:15.

14. Morris R, Lord S, Lawson RA, et al. Gait Rather Than Cognition Predicts Decline in Specific Cognitive Domains in Early Parkinson’s Disease. J Gerontol A Biol Sci Med Sci 2017;72:1656–1662.

15. Giladi N, Huber-Mahlin V, Herman T, Hausdorff JM. Freezing of gait in older adults with high level gait disorders: association with impaired executive function. J Neural Transm (Vienna) 2007;114:1349–1353.

16. Amboni M, Cozzolino A, Longo K, Picillo M, Barone P. Freezing of gait and executive functions in patients with Parkinson’s disease. Mov Disord 2008;23:395–400.

17. Rushworth MF, Hadland KA, Paus T, Sipila PK. Role of the human medial frontal cortex in task switching: a combined fMRI and TMS study. J Neurophysiol 2002;87:2577–2592.

18. Ragozzino ME. The contribution of the medial prefrontal cortex, orbitofrontal cortex, and dorsomedial striatum to behavioral flexibility. Ann N Y Acad Sci 2007;1121:355–375.

19. Playford ED, Jenkins IH, Passingham RE, Nutt J, Frackowiak RS, Brooks DJ. Impaired mesial frontal and putamen activation in Parkinson’s disease: a positron emission tomography study. Ann Neurol 1992;32:151–161.

20. Singh A, Richardson SP, Narayanan N, Cavanagh JF. Mid-frontal theta activity is diminished during cognitive control in Parkinson’s disease. Neuropsychologia 2018;117:113–122.

21. Toledo JB, Lopez-Azcarate J, Garcia-Garcia D, et al. High beta activity in the subthalamic nucleus and freezing of gait in Parkinson’s disease. Neurobiol Dis 2014;64:60–65.

22. Wagner J, Makeig S, Gola M, Neuper C, Muller-Putz G. Distinct beta Band Oscillatory Networks Subserving Motor and Cognitive Control during Gait Adaptation. J Neurosci 2016;36:2212–2226.

23. Singh A. Oscillatory activity in the cortico-basal ganglia-thalamic neural circuits in Parkinson’s disease. Eur J Neurosci 2018;48:2869–2878.

24. Singh A, Plate A, Kammermeier S, Mehrkens JH, Ilmberger J, Bötzel K. Freezing of gait-related oscillatory activity in the human subthalamic nucleus. Basal Ganglia 2013;3:25–32.

25. Kelley R, Flouty O, Emmons EB, et al. A human prefrontal-subthalamic circuit for cognitive control. Brain 2018;141:205–216.

26. Shine JM, Handojoseno AM, Nguyen TN, et al. Abnormal patterns of theta frequency oscillations during the temporal evolution of freezing of gait in Parkinson’s disease. Clin Neurophysiol 2014;125:569–576.

27. Nutt JG, Bloem BR, Giladi N, Hallett M, Horak FB, Nieuwboer A. Freezing of gait: moving forward on a mysterious clinical phenomenon. Lancet Neurol 2011;10:734–744.

28. Cavanagh JF, Frank MJ. Frontal theta as a mechanism for cognitive control. Trends Cogn Sci 2014;18:414–421.

29. Sharott A, Magill PJ, Harnack D, Kupsch A, Meissner W, Brown P. Dopamine depletion increases the power and coherence of beta-oscillations in the cerebral cortex and subthalamic nucleus of the awake rat. Eur J Neurosci 2005;21:1413–1422.

30. Anidi C, O’Day JJ, Anderson RW, et al. Neuromodulation targets pathological not physiological beta bursts during gait in Parkinson’s disease. Neurobiol Dis 2018;120:107–117.

31. Gibb WR, Lees AJ. A comparison of clinical and pathological features of young- and old-onset Parkinson’s disease. Neurology 1988;38:1402–1406.

32. Giladi N, Shabtai H, Simon ES, Biran S, Tal J, Korczyn AD. Construction of freezing of gait questionnaire for patients with Parkinsonism. Parkinsonism Relat Disord 2000;6:165–170.

33. Nasreddine ZS, Phillips NA, Bedirian V, et al. The Montreal Cognitive Assessment, MoCA: a brief screening tool for mild cognitive impairment. J Am Geriatr Soc 2005;53:695–699.

34. Movement Disorder Society Task Force on Rating Scales for Parkinson’s D. The Unified Parkinson’s Disease Rating Scale (UPDRS): status and recommendations. Mov Disord 2003;18:738–750.

35. Vercruysse S, Gilat M, Shine JM, Heremans E, Lewis S, Nieuwboer A. Freezing beyond gait in Parkinson’s disease: a review of current neurobehavioral evidence. Neurosci Biobehav Rev 2014;43:213–227.

36. Abe K, Asai Y, Matsuo Y, et al. Classifying lower limb dynamics in Parkinson’s disease. Brain Res Bull 2003;61:219–226.

37. Nolan H, Whelan R, Reilly RB. FASTER: Fully Automated Statistical Thresholding for EEG artifact Rejection. J Neurosci Methods 2010;192:152–162.

38. Cohen MX. Analyzing Neural Time Series Data: Theory and Practice (Issues in Clinical and Cognitive Neuropsychology): The MIT Press, 2014.

39. Morris R, Lord S, Bunce J, Burn D, Rochester L. Gait and cognition: Mapping the global and discrete relationships in ageing and neurodegenerative disease. Neurosci Biobehav Rev 2016;64:326–345.

40. Kelly VE, Eusterbrock AJ, Shumway-Cook A. A review of dual-task walking deficits in people with Parkinson’s disease: motor and cognitive contributions, mechanisms, and clinical implications. Parkinsons Dis 2012;2012:918719.

41. Hausdorff JM, Balash J, Giladi N. Effects of cognitive challenge on gait variability in patients with Parkinson’s disease. J Geriatr Psychiatry Neurol 2003;16:53–58.

42. Wager TD, Jonides J, Reading S. Neuroimaging studies of shifting attention: a meta-analysis. Neuroimage 2004;22:1679–1693.

43. Rosenberg-Katz K, Herman T, Jacob Y, Giladi N, Hendler T, Hausdorff JM. Gray matter atrophy distinguishes between Parkinson disease motor subtypes. Neurology 2013;80:1476–1484.

44. Criado JM, de la Fuente A, Heredia M, Riolobos AS, Yajeya J. Electrophysiological study of prefrontal neurones of cats during a motor task. Pflugers Arch 1997;434:91–96.

45. Walton CC, Shine JM, Mowszowski L, Naismith SL, Lewis SJ. Freezing of gait in Parkinson’s disease: current treatments and the potential role for cognitive training. Restor Neurol Neurosci 2014;32:411–422.

46. Meissner SN, Krause V, Sudmeyer M, Hartmann CJ, Pollok B. The significance of brain oscillations in motor sequence learning: Insights from Parkinson’s disease. Neuroimage Clin 2018;20:448–457.

47. Singh A, Trapp NT, De Corte B, et al. Cerebellar Theta Frequency Transcranial Pulsed Stimulation Increases Frontal Theta Oscillations in Patients with Schizophrenia. Cerebellum 2019;18:489–499.

48. Parker KL, Chen KH, Kingyon JR, Cavanagh JF, Narayanan NS. Medial frontal approximately 4-Hz activity in humans and rodents is attenuated in PD patients and in rodents with cortical dopamine depletion. J Neurophysiol 2015;114:1310–1320.

49. Caplan JB, Madsen JR, Schulze-Bonhage A, Aschenbrenner-Scheibe R, Newman EL, Kahana MJ. Human theta oscillations related to sensorimotor integration and spatial learning. J Neurosci 2003;23:4726–4736.

50. Miller EK, Cohen JD. An integrative theory of prefrontal cortex function. Annu Rev Neurosci 2001;24:167–202.

51. Singh A, Kammermeier S, Mehrkens JH, Botzel K. Movement kinematic after deep brain stimulation associated microlesions. J Neurol Neurosurg Psychiatry 2012;83:1022–1026.

52. Singh A, Mehrkens JH, Botzel K. Effect of micro lesions of the basal ganglia on ballistic movements in patients with deep brain stimulation. J Neurol Sci 2012;314:175–177.

53. Morris ME. Movement disorders in people with Parkinson disease: a model for physical therapy. Phys Ther 2000;80:578–597.

54. Vercruysse S, Spildooren J, Heremans E, et al. Freezing in Parkinson’s disease: a spatiotemporal motor disorder beyond gait. Mov Disord 2012;27:254–263.

55. Delval A, Moreau C, Bleuse S, et al. Auditory cueing of gait initiation in Parkinson’s disease patients with freezing of gait. Clin Neurophysiol 2014;125:1675–1681.

56. Donovan S, Lim C, Diaz N, et al. Laserlight cues for gait freezing in Parkinson’s disease: an open-label study. Parkinsonism Relat Disord 2011;17:240–245.

57. Lee SJ, Yoo JY, Ryu JS, Park HK, Chung SJ. The effects of visual and auditory cues on freezing of gait in patients with Parkinson disease. Am J Phys Med Rehabil 2012;91:2–11.

58. Vercruysse S, Devos H, Munks L, et al. Explaining freezing of gait in Parkinson’s disease: motor and cognitive determinants. Mov Disord 2012;27:1644–1651.

59. Cools R, Miyakawa A, Sheridan M, D’Esposito M. Enhanced frontal function in Parkinson’s disease. Brain 2010;133:225–233.

60. Cools R, D’Esposito M. Inverted-U-shaped dopamine actions on human working memory and cognitive control. Biol Psychiatry 2011;69:e113–125.

61. Hanakawa T, Katsumi Y, Fukuyama H, et al. Mechanisms underlying gait disturbance in Parkinson’s disease: a single photon emission computed tomography study. Brain 1999;122 (Pt 7):1271–1282.

62. Shine JM, Matar E, Ward PB, et al. Freezing of gait in Parkinson’s disease is associated with functional decoupling between the cognitive control network and the basal ganglia. Brain 2013;136:3671–3681.

